# RELATIONSHIP BETWEEN PARENTAL AGE AND SEVERITY OF OROFACIAL CLEFTS

**DOI:** 10.1101/2021.02.22.432246

**Authors:** Olawale Adamson, Abimibola V. Oladugba, Azeez Alade, Waheed O. Awotoye, Tamara Busch, Mary Li, Joy Olotu, Veronica C. Sule, Azeez Fashina, James Olutayo, Mobolanle O. Ogunlewe, Wasiu L. Adeyemo, Azeez Butali

## Abstract

**OBJECTIVES:** This study aims to investigate the relationship between paternal age, maternal age, and both on the severity of orofacial clefts.

**DESIGN:** This was a retrospective study of cases which were subjects clinically diagnosed with non-syndromic cleft lip and/or palate (CL/P). Data was obtained from the AFRICRAN project database on Nigerian non-syndromic orofacial cleft cases.

**SETTING:** The samples for cases in this study were obtained at the Cleft clinic of Oral and Maxillofacial surgery at the Lagos University Teaching Hospital, Lagos.

**OUTCOME:** Primary outcome measure is severity of orofacial clefts and secondary outcome measure is to evaluate the effect of parental age in determining the incidence of left or right sided orofacial clefts.

**RESULTS:** There is no statistical significant association between type of CL ± P and parental age in young fathers (p=0.93). When old fathers are considered, percentage of complete (more severe) CL ± P cases increases especially in old mothers and this was statistically significant (p=0.036). In old fathers, the risk of CL ± P is increased (OR: 2.66, CI: 1.04-6.80) and also there is increased risk of developing right sided CL ± P (OR: 1.61, CI: 1.0-2.59). There is reduced risk of isolated cleft palate in young fathers (OR: 0.36, CI: 0.07-1.71) but the risk increases when considering complete types (more severe) of isolated cleft palates (OR: 1.63, CI: 0.71-3.7)

**CONCLUSION:** The study shows a higher risk of CL ± P is associated with increase father’s age.

## INTRODUCTION

Birth defects are reported to contribute significantly to infant morbidity and mortality globally (1). Orofacial clefts (OFC) are amongst the most common craniofacial birth defects with a prevalence of 1:700 live births (2). OFC can be syndromic or non-syndromic with syndromic accounting for 70% of all OFC (Dixon et al., 2011). The phenotypic presentation of OFC differs and ranges from cleft lip (CL), cleft lip and palate (CLP) and cleft palate only (CPO). The aetiology of OFC is considered to be multifactorial with polygenic, environmental, epigenetics and interaction between genetics and environmental factors playing a role (3). Environmental factors implicated in aetiology of OFC include smoking, alcohol, metabolic syndromes such as diabetes mellitus and maternal obesity as well as parental age.

Parental age has been proposed as a possible risk factor for OFC.(4) Previous studies conducted on the association between parental age and incidence of birth defects have yielded inconsistent results(5)(6,7). It is generally reported that advanced age may predispose chromosomes to irreversible changes and genetic alterations. In a study by Sartorelli et al.,(8) the frequency of numerical and structural chromosomal aberrations (acentric fragments and complex radial figures) was significantly greater in chromosomes of older donors when compared with those of the younger group. Many autosomal dominant diseases have been shown to be associated with increasing paternal age.(9) Crouzon syndrome, Apert syndrome and Pfeiffer syndrome are all autosomal dominant craniosynostosis disorders that can be caused by mutations in the FGFR2 gene occurring in a normal father’s germ line. All the FGFR2 mutations were associated with increased paternal age and molecularly proven to be of paternal origin.(10) A Danish population-based study of 1,920 OFC affected births of 1,489,014 live births concluded that paternal age is associated with CLP, independently of maternal age.(11). It is worthy to note that the fetal congenital anomalies attributed to advanced paternal age is low in absolute terms and though there is a relationship, it is not causal in effect.(9)

There are studies suggesting that maternal age and parity might play an important role in the development of certain isolated birth defects.(12). Kim et al (13) reported that risk of trisomy 21, trisomy 18, triple X syndrome, and all aneuploidies showed a significant increase related to increase in maternal age. For Down syndrome, the risk of maternal age did not change when controlling for paternal age. On the other hand, paternal age effects changed from very large risk to a small sparing risk when controlling for maternal age.

There is no clear consensus on the effect of parental age regarding the risk of orofacial clefts though many studies have reported associations between advanced maternal or paternal age and risk of orofacial clefts. A study by Bille et al(4) using the population-based Danish Facial Cleft Database, reported that the influence of maternal and paternal ages on the risk of Cleft lip and/or palate (CL ± P) increases with the advancing age of the other parent, and that the influence vanishes if the other parent is young. In contrast, the risk of having a child with cleft palate is influenced only by father’s age, not mother’s age. In a study of Brazilians with OFC, Martelli et al(14) reported an association between maternal age and increased risk for CLP while paternal age risk is not significant.

In addition to the fact that the association between parental age and risk of orofacial clefts has been inconsistent, there is sparse literature on the influence of parental age on the severity of orofacial clefts. This study aims to investigate the relationship between paternal age, maternal age, and both on the severity of orofacial clefts. In addition, we plan to evaluate the effect of parental age in determining the incidence of left or right sided orofacial clefts.

## MATERIAL AND METHODS

### Study design

This was a retrospective study in which cases were subjects clinically diagnosed with non-syndromic cleft lip and/or palate (CL ± P). Selection of cases was based on standardized examination performed by trained Surgeons who participated in the Pan-African Association of Cleft lip and palate network for repair of orofacial clefts in Africa. Clinical information including detailed description of the phenotype, parental age and clinical photographs were recorded in the database.

### Study location

The samples for cases in this study were obtained at the Cleft clinic of Oral and Maxillofacial surgery at the Lagos University Teaching Hospital, Lagos. The Research and ethics committee of Lagos University Teaching Hospital was informed, and ethical approval obtained before commencing the study.

### Method

Data was obtained from the AFRICRAN project database on Nigerian non-syndromic orofacial cleft cases. All infants born with orofacial clefts were clinically examined with the overall goal to measure and characterize the craniofacial morphology and development, and data on parental age were also included. The infants were classified according to whether they were unilateral (left [L] or right [R] sided) or bilateral, as well as the severity of their cleft graded.

For the current analysis regarding the influence of parental age on cleft severity, the groups with CL ± P and isolated cleft palate were considered two separate populations because of their different embryological origins. The CL ± P population comprised: unilateral incomplete cleft lip (UICL), bilateral incomplete cleft lip (BICL), unilateral complete cleft lip (UCCL), bilateral complete cleft lip (BCCL), unilateral incomplete cleft lip and palate (UICLP), bilateral incomplete cleft lip and palate (BICLP), unilateral complete cleft lip and palate (UCCLP) and BCCLP (bilateral complete cleft lip and palate).

In the CL ± P population, the data were grouped by the analysis of the influence of severity. For this purpose, the previously described subgroups were combined as follows: IC (incomplete/less severe clefts = UICL+BICL+UICLP+BICLP) vs CC (complete/ more severe clefts = UCCL+BCCL+UCCLP+BCCLP), as well as L vs R-sided cleft (for this analysis, only unilateral clefts (UCL ± P = UICL+UCCL+UICLP+UCCLP) were included).

The CP population comprised of: incomplete cleft palate (ICP) and complete cleft palate (CCP)

The parental age was classified into young father, old father or young mother, old mother based on the median ages of the parents. The risk of orofacial clefts was analyzed based on these groups.

Statistical analysis: For the primary analysis, a binary outcome variable was defined with two values (0 = IC, 1 = CC). Pearson’s Chi-square test was applied to analyze the association between parental age and the severity of orofacial clefts. Based on logistic regression, the relative risk with confidence interval was calculated between severity of orofacial clefts and parental age.

## RESULTS

The total number of non-syndromic orofacial cleft cases analyzed was 267 with 202 CL ± P and 65 CP cases. Table 1 shows the parental age distribution of the cleft cases. Young fathers are categorized as those below 35 years while old fathers are greater than or equals to 35 years old while young mothers are categorized as those below 30 years while old mothers are greater than or equals to 30 years old.

**Table 1:**
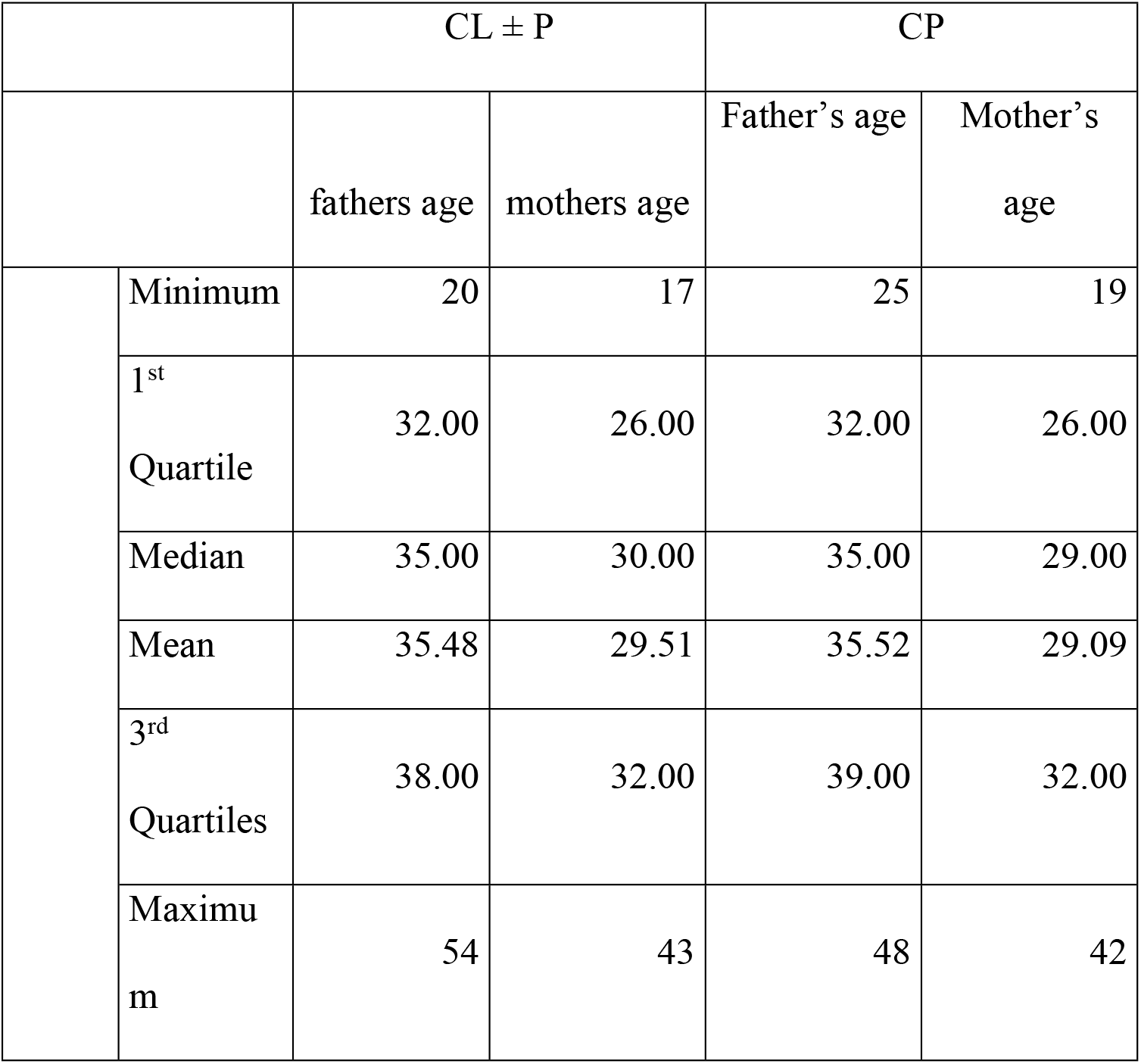
Summary statistics of the age distribution of the father and mother in the data.

| | | CL ± P | | CP | |
| --- | --- | --- | --- | --- | --- |
|  |  | fathers age | mothers age | Father's age | Mother's age |
|  | Minimum | 20 | 17 | 25 | 19 |
|  | 1<sup>st</sup><br>Quartile | 32.00 | 26.00 | 32.00 | 26.00 |
|  | Median | 35.00 | 30.00 | 35.00 | 29.00 |
|  | Mean | 35.48 | 29.51 | 35.52 | 29.09 |
|  | 3<sup>rd</sup><br>Quartiles | 38.00 | 32.00 | 39.00 | 32.00 |
|  | Maximum | 54 | 43 | 48 | 42 |

### CLEFT LIP AND PALATE

Generally, there are more complete cleft CL ± P than incomplete cases (Table 2). There is no statistical significant association between type of CL ± P and parental age in young fathers (p=0.93). When old fathers are considered, percentage of complete CL ± P cases increases especially in old mothers and this was statistically significant at (p=0.036). These findings indicate that old father-old mother combination is more associated with more severe CL ± P.

**Table 2:** Shows the relationship of parental age to cleft lip and palate cases.

| Fathers age based on median |  |  |  | Type of Cleft lip |  | Total | p-value |
| --- | --- | --- | --- | --- | --- | --- | --- |
|  |  |  |  | IC | CC |  |  |
| < 35 | Mothers age based on median | < 30 | Count | 13 | 42 | 55 | 0.930 |
|  |  |  | % of Total | 16.9% | 54.5% | 71.4% |  |
|  |  | > 30 | Count | 5 | 17 | 22 |  |
|  |  |  | % of Total | 6.5% | 22.1% | 28.6% |  |
|  | Total |  | Count | 18 | 59 | 77 |  |
|  |  |  | % of Total | 23.4% | 76.6% | 100.0% |  |
| ≥35 | Mothers age based on median | < 30 | Count | 12 | 32 | 44 | 0.036 |
|  |  |  | % of Total | 9.6% | 25.6% | 35.2% |  |
|  |  | > 30 | Count | 10 | 71 | 81 |  |
|  |  |  | % of Total | 8.0% | 56.8% | 64.8% |  |
|  | Total |  | Count | 22 | 103 | 125 |  |
|  |  |  | % of Total | 17.6% | 82.4% | 100.0% |  |

#### Severity of CL ± P

There is no increased risk of CL ± P in young fathers (OR: 1.05, CI: 0.3-3.4) and there is no increased risk for any subtype of CL ± P (table 3). In old fathers, the risk of CL ± P is increased (OR: 2.66, CI: 1.04-6.80). There is also increased risk for incomplete CL ± P in old fathers (OR: 2.209, CI: 1.04-4.70) but the risk reduces when complete CL ± P was considered (OR: 0.83, CI: 0.68-1.01). These show that the risk of CL ± P increases with paternal age which is higher in less severe form of incomplete CL ± P (Table 3).

**Table 3:** Relative risk of severity of CL ± P in relation to parental age.

| Risk Estimate |  |  |  |  |
| --- | --- | --- | --- | --- |
| Fathers age based on median |  | Value | 95% Confidence Interval |  |
|  |  |  | Lower | Upper |
| < 35 | Odds Ratio for Mothers age based on median (< 30 / > 30) | 1.052 | .325 | 3.409 |
|  | For cohort Type of Cleft lip = IC | 1.040 | .421 | 2.571 |
|  | For cohort Type of Cleft lip = CC | .988 | .754 | 1.295 |
| ≥ 35 | Odds Ratio for Mothers age based on median (< 30 / > 30) | 2.663 | 1.043 | 6.797 |
|  | For cohort Type of Cleft lip = IC | 2.209 | 1.039 | 4.699 |
|  | For cohort Type of Cleft lip = CC | .830 | .680 | 1.012 |
| Total | Odds Ratio for Mothers age based on median (< 30 / > 30) | 1.982 | .974 | 4.035 |
|  | For cohort Type of Cleft lip = IC | 1.734 | .973 | 3.090 |
|  | For cohort Type of Cleft lip = CC | .875 | .761 | 1.006 |

#### Risk of left or right sided cleft in unilateral CL ± P

There is no associated increase in risk of unilateral CL ± P for either left or right side in young fathers (Table 4). In old fathers, there is increased risk of developing right sided CL ± P (OR: 1.61, CI: 1.0-2.59) and the risk of developing left sided clefts reduces indicating that mother’s age is a more associated with left-sided clefts in old fathers.

**Table 4:** Relative risk of Left or right sided Unilateral CL ± P.

| Risk Estimate |  |  |  |  |
| --- | --- | --- | --- | --- |
| Fathers age based on median |  | Value | 95% Confidence Interval |  |
|  |  |  | Lower | Upper |
| < 35 | Odds Ratio for Mothers age based on median (< 30 / > 30) | 1.045 | .352 | 3.104 |
|  | For cohort Cleft Lip Details = Left | 1.026 | .548 | 1.919 |
|  | For cohort Cleft Lip Details = Right | .981 | .618 | 1.558 |
| ≥ 35 | Odds Ratio for Mothers age based on median (< 30 / > 30) | .442 | .193 | 1.011 |
|  | For cohort Cleft Lip Details = Left | .714 | .493 | 1.033 |
|  | For cohort Cleft Lip Details = Right | 1.614 | 1.006 | 2.588 |
| Total | Odds Ratio for Mothers age based on median (< 30 / > 30) | .508 | .273 | .943 |
|  | For cohort Cleft Lip Details = Left | .733 | .549 | .978 |
|  | For cohort Cleft Lip Details = Right | 1.443 | 1.028 | 2.025 |

#### Severity of bilateral CL ± P

In bilateral CL ± P, there is a slight risk of bilateral CL ± P in young fathers (OR: 1.14, CI: 0.7-16.94) (Table 5). There was two-fold increase in risk of bilateral CL ± P in old fathers (OR: 2.0, CI: 0.11-36.9) and this was more predominant in incomplete bilateral CL ± P (OR: 1.87, CI: 0.13-26.1).

**Table 5:** Relative risk of bilateral CL ± P.

| Risk Estimate |  |  |  |  |
| --- | --- | --- | --- | --- |
| Fathers age based on median |  | Value | 95% Confidence Interval |  |
|  |  |  | Lower | Upper |
| < 35 | Odds Ratio for Mothers age based on median (< 30 / > 30) | 1.143 | .077 | 16.947 |
|  | For cohort Type of Cleft lip = IC | 1.091 | .184 | 6.476 |
|  | For cohort Type of Cleft lip = CC | .955 | .382 | 2.387 |
| ≥ 35 | Odds Ratio for Mothers age based on median (< 30 / > 30) | 2.000 | .108 | 36.954 |
|  | For cohort Type of Cleft lip = IC | 1.875 | .134 | 26.161 |
|  | For cohort Type of Cleft lip = CC | .938 | .698 | 1.259 |
| Total | Odds Ratio for Mothers age based on median (< 30 / > 30) | 2.857 | .477 | 17.110 |
|  | For cohort Type of Cleft lip = IC | 2.368 | .524 | 10.698 |
|  | For cohort Type of Cleft lip = CC | .829 | .605 | 1.135 |

### CLEFT PALATE

There is reduced risk of isolated cleft palate in young fathers (OR: 0.36, CI: 0.07-1.71) but the risk increases when considering complete cleft palates (OR: 1.63, CI: 0.71-3.7) though this was not statistically significant (Table 6). This indicates that maternal age is more associated with less severe cleft palate while paternal age is associated with more severe cleft palate.

**Table 6:** Risk of parental age and severity of cleft palate only.

| Risk Estimate |  |  |  |  |
| --- | --- | --- | --- | --- |
| Fathers age based on median |  | Value | 95% Confidence Interval |  |
|  |  |  | Lower | Upper |
| <35 | Odds Ratio for Mothers age based on median (<30 / >30) | .359 | .075 | 1.714 |
|  | For cohort Populations = ICP | .583 | .267 | 1.276 |
|  | For cohort Populations = CCP | 1.625 | .713 | 3.706 |
| ≥35 | Odds Ratio for Mothers age based on median (<30 / >30) | .593 | .149 | 2.365 |
|  | For cohort Populations = ICP | .781 | .395 | 1.545 |
|  | For cohort Populations = CCP | 1.316 | .646 | 2.680 |
| Total | Odds Ratio for Mothers age based on median (<30 / >30) | .445 | .165 | 1.200 |
|  | For cohort Populations = ICP | .663 | .398 | 1.106 |
|  | For cohort Populations = CCP | 1.492 | .904 | 2.462 |

## DISCUSSION

This study evaluates the relationship between parental age and severity of cleft using data derived from Nigerian patients with cleft lip and palate. To our knowledge, this is the first of such study to be conducted in an African population. This study attempts to go further than just linking parental age with risk of OFC, but highlighting the effects on severity and also on left or right selection of unilateral cleft lip and palate cases.

This study shows that increased parental age is associated with more severe CL ± P cases as combination of older parents produce more severe cases. This aligns with various studies that have reported increased congenital malformations in older parents.(6,15). A population-based study on Danish Facial Cleft Database, reported the influence of maternal and paternal ages on the risk of cleft lip with or without cleft palate increases with the advancing age of the other parent, and that the influence vanishes if the other parent is young.(7) Though there has been varying reports on whether the maternal or paternal chromosomes are culpable. The exact mechanism of this occurrence has not been elucidated, though single gene mutations are suggested mechanisms.(11)

Also from this study, advanced paternal age is associated with increased risk of less severe unilateral and bilateral CL ± P. This is in agreement with a similar study by Herman et al(16) that reported that paternal age increases risk of CL ± P which is more pronounced with advanced maternal age. The paternal age seems to have a great deal of influence on the prevalence of CL ± P in any population. The influence of paternal age also spills over in to cleft palate where many studies have reported association between paternal age and cleft palate.(Martelli *et al.*, 2010; Hoda Badr *et al.*, 2011) In this study, paternal age is associated with increased risk of more severe cleft palate.

Though maternal age has been associated with chromosomal abnormalities in some studies but paternal age is usually associated with birth defects.(12,13) It is reported in some literature that the risk of birth defects such as heart malformation, other musculoskeletal anomalies, tracheo-oesophageal fistula/oesophageal atresia, Down’s syndrome and other chromosomal anomalies, increases slightly with advancing paternal age.(5,14). Association between younger fathers and several selected birth defects like neural tube defects has also been published (18) The association of paternal age with birth defects has been attributed to accumulation of chromosomal aberrations and mutations during the maturation of male germ cells.(10,19). The amount of DNA damage in sperm of men aged 36–57 is three times that of men 35 years and less.(11)

Prevalence and pattern of occurrence of OFC in a given population is expected to fluctuate as the average parental ages change. Increase occurrence of more severe cleft is expected with advanced parental ages and this may take a toll on available resources.

### Strengths and Limitations

The strength of this study is that it is a population-based investigation of a genetically homogeneous population who has similar environmental exposures. Furthermore, only parents of children with non-syndromic cleft were included. The limitation of this study is the small sample size and other environmental factors like socio-economic status of the parents, maternal intake of alcohol and smoking were not considered

### Conclusion

Increased parental age is associated with increased risk of OFC. In this study, advanced paternal age is associated with increased risk of less severe unilateral and bilateral CL ± P but a more severe cleft palate. Future prospective studies on different populations and also considering other socio-economic factors may provide more insights into the influence of parental age on occurrence and severity of OFC.

## ACKNOWLEDGEMENTS

We are grateful to the families who voluntarily participated in this study in Nigeria. We are also grateful to all the administrative and research staffs, students, nurses and resident doctors who assisted with participant recruitment, consent, and data collection. This research is supported by the National Institute of Dental and Craniofacial Research (R00 DE022378 and R01DE028300; A.B.).

